# Animal virus ecology and evolution are shaped by the virus host-body infiltration and colonization pattern

**DOI:** 10.1101/492603

**Authors:** Jan Slingenbergh

**Affiliations:** Former FAO of the UN. Current address: Schuettorf, Niedersachsen, Germany

## Abstract

The current classification of animal viruses primarily relates to the virus molecular world, the genomic architecture and the corresponding host-cell infection cycle. This virus centered perspective does not make allowance for the precept that virus fitness hinges on the virus transmission success. Virus transmission reflects the infection-shedding-transmission dynamics and, with it, the organ system involvement and other, macroscopic dimensions of the host environment. This study examines the transmission ecology of the world main livestock viruses, 36 in total, belonging to eleven different families, and a mix of RNA, DNA and retroviruses. Viruses are virtually ranked in an outer- to inner-body fashion, based on the shifting organ system involvement and associated infection-shedding-transmission dynamics. As a next step, this ranking is disentangled with the aim to contrast two main host ecologies, poultry plus pig production and ruminant plus equine husbandry, as well as to create a distinction among the RNA, DNA and retroviruses, also ranked in an outer- to inner-body fashion. Spearman correlation reveals the matches among these various virus traits, as pertaining to the two host-ecologies, four infection-shedding-transmission related variables, and the three virus genomes. The collective results reveal the outer- to inner-body shifts in the interplay of host environment, virus-host interactions, and nature of the virus. Two opposing virus evolution pathways emerge, respectively for generalist type, outer-body and for specialist type, inner-body viruses. The ecological virus classification here presented is broadly consistent with the current virus classification system and offers the advantage of bringing substance and cohesion to the interrelationships among viruses and virus families.

**Author Summary:** It remains unknown how exactly viruses fit in the tree of life. Still, there is growing awareness that viruses as biological replicators are subjected to ecological sorting and so require a viable propagation strategy. In the current analysis I depart from the precept that virus fitness hinges on the virus transmission success. I examine the transmission ecologies of the world main livestock viruses, 36 in total, a collection of pathogens well described in terms of the organ system involvement, infection course, the extent of host damage, virus shedding profile, and virus transmission modes. The viruses are on this basis ranked in an outer- to inner-body fashion, virtually. As a next step, this ranking is disentangled with a view to contrast two main host ecologies, poultry plus pig production and ruminant plus equine husbandry, as well as to create a distinction among the RNA, DNA and retrovirus in the study. The matches among these various virus traits serve to establish the outer- to inner-body shifts in the interplay of host environment, virus-host interactions, and nature of the virus. Two opposing virus evolution pathways emerge, respectively for generalist type, outer-body and for specialist, inner-body viruses.

## Introduction

Increasingly, viruses are viewed in an ecological context, as living entities [1,2,3,4,5]. Viruses, like all biological replicators, form a continuum along the selfishness-cooperativity axis, from the completely selfish to fully cooperative forms [6]. In a deeper sense, the history of life is a story of parasite-host coevolution that includes both the incessant arms race and various forms of cooperation [7].

### Organ systems and virus transmission success

The current paper approaches the evolution of viral replication and propagation strategies from a practical angle and departs from the virus transmission success. Virus fitness hinges on the ability of the virus to transmit to other hosts. There are macroscopic dimensions to virus transmission. For example, a virus may establish in the upper respiratory tract and transmit by air via aerosols. An enteric virus features a fecal-oral cycle. A skin virus may transmit on the basis of touch or abrasion. A virus colonizing the distal urogenital tract and the external genitalia may transmit during sexual contact. Hence, the analysis of animal virus transmission requires consideration of the overt clinical signs that associate with the infection process, the gross pathology, and of how the infection translates in a virus specific shedding profile.

### Outer-versus inner-body and macroscopic versus molecular level

The virus organ system tropism may be assumed to evolve in harmony with the virus cell tropism and so links the macroscopic to the molecular level virus-host interactions. This may be further illustrated with the dichotomy in the release of viruses from epithelial cells. Apart from direct cell-to-cell transmission [8], viruses may be released from the apical cell surface and so end-up in the outer-body environment. To make it to the next host, these viruses colonize the various mucosae or also the skin. Conversely, viruses released from the basolateral cell surface infiltrate underlying tissues. These viruses may end-up in the lymph drainage and from there enter the blood circulation and so infect any of the internal organs. Infiltration of and establishment in the inner-body environment may translate in a number of additional, non-epithelial transmission modes. For example, virus establishment in the reproductive organs may yield seminal, trans-placental, congenital, or also lactogenic transmission. In birds, virus may be shed into yolk or albumen and so transmit vertically [9]. Virus circulation in the bloodstream may result in virus transmission via arthropod vectors or needles [10]. A rabies virus replicates in the central nervous system and so enhances virus transmission [11].

### The study

The study explores the transmission ecology of the world main livestock viruses, 36 in total, a collection of well described pathogens. Viruses are ranked in an outer- to inner-body fashion, virtually, on the basis of the shifting organ system involvement, transmission modes, and infection-shedding-transmission related variables. Next, this virus infiltration ranking is disentangled to contrast the poultry plus pig to the ruminant plus equine viruses, as well as to separately consider the RNA, DNA and retroviruses in the study, also ranked in an outer- to inner-body fashion. The various virus traits, as pertaining to the host-ecology, infection-shedding-transmission related variables, and virus genome, are all matched on the basis of Spearman correlation. A synthesis of these findings reveals the outer- to inner-body shifts in the interplay of host ecology, virus-host interactions, and nature of the virus. What results is a predictive framework for animal virus evolution.

## Results

### Matching virus ecological variables

As a first step, the one-to-three scores allocated to the 36 livestock viruses for the four ecological variables were matched on the basis of Spearman correlation, see also S2 Fig. The extent of virus host-body infiltration and the length of the infection period were found correlated, with R = 0.78 and P = 0. In addition, the extent of virus infiltration and the virus environmental survival rate were found correlated, negatively, with R = - 0.44 and P < 0.01.

### A virus family specific infiltration ranking

Next, the eleven virus families in the study were grouped and ranked A-D on the basis of the infiltration scores allocated to the family viruses, see Fig 1. The transmission of the viruses belonging to the *Orthomyxoviridae* and the *Paramyxoviridae* families was found to result strictly from epithelial infections. Next, the transmission of the viruses belonging to the *Coronaviridae*, the *Picornaviridae* and the *Poxviridae* was in part found modulated by internal organ systems. Next, the transmission of the viruses belonging to the *Arteriviridae*, the *Flaviviridae*, the *Herpesviridae*, plus also the single infectious bursal disease virus, was found modulated by both epithelia and internal organ systems. Finally, the transmission of the viruses belonging to the *Retroviridae* family plus the single bluetongue virus was found modulated by epithelia and internal organ systems or by just internal organ systems. Spearman correlation based on the new, family specific, one-to-four virus host-body infiltration score and the length of the infection period score yielded an R = 0.86 and P = 0. The new match with the virus environmental survival rate score yielded an R = - 0.46 and P < 0.005. When virus environmental survival was considered just for the outer-body viruses, virus infiltration and environmental survival were found positively correlated, with R = 0.53 and P < 0.05. For the inner-body viruses, infiltration and virus environmental survival were negatively correlated, with R = - 0.75 and P < 0.0005. In addition, virus infiltration and infection severity were found correlated, negatively, with R = - 0.38 and P < 0.05.

**Fig 1.**
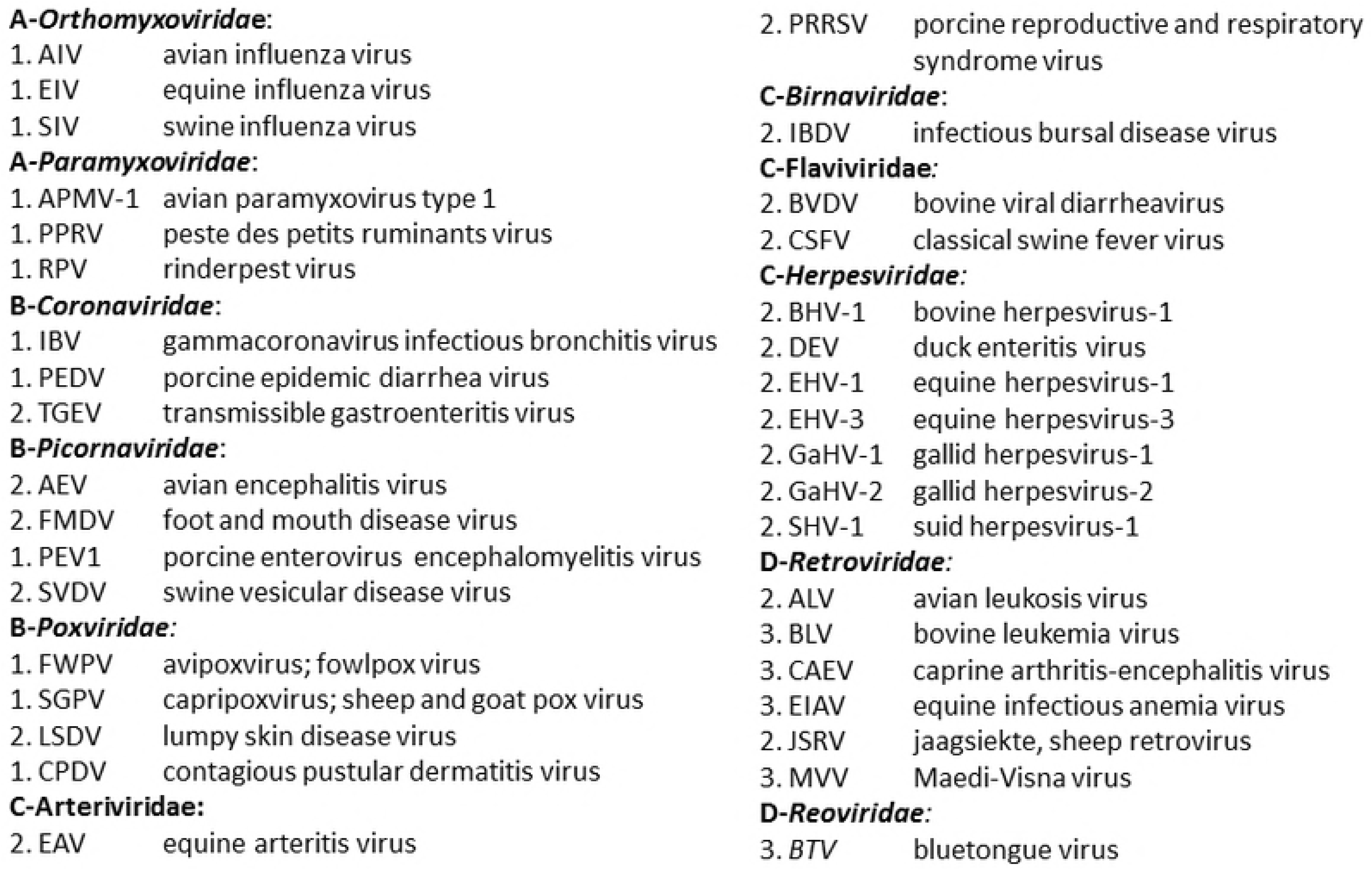
An outer- to inner-body line-up of virus family groupings. The eleven virus families in the study were grouped and ranked A-D on the basis of the host-body infiltration scores allocated to the family viruses. For each grouping, the virus families and also the family viruses are shown in alphabetical order.

### A virtual, outer- to inner-body line-up of organ systems

Next, the organ system tropisms of the viruses belonging to each family were collectively fitted to and with the naked eye aligned with the Fig 1 family group line-up. As indicated in Fig 2, the resulting, virtual outer- to inner-body line-up of organ systems runs from the respiratory plus alimentary tracts to the skin, distal urogenital tract or cloaca, peripheral nerves and ganglia, reproductive organs system, lungs, to the immune and circulatory systems.

**Fig 2.**
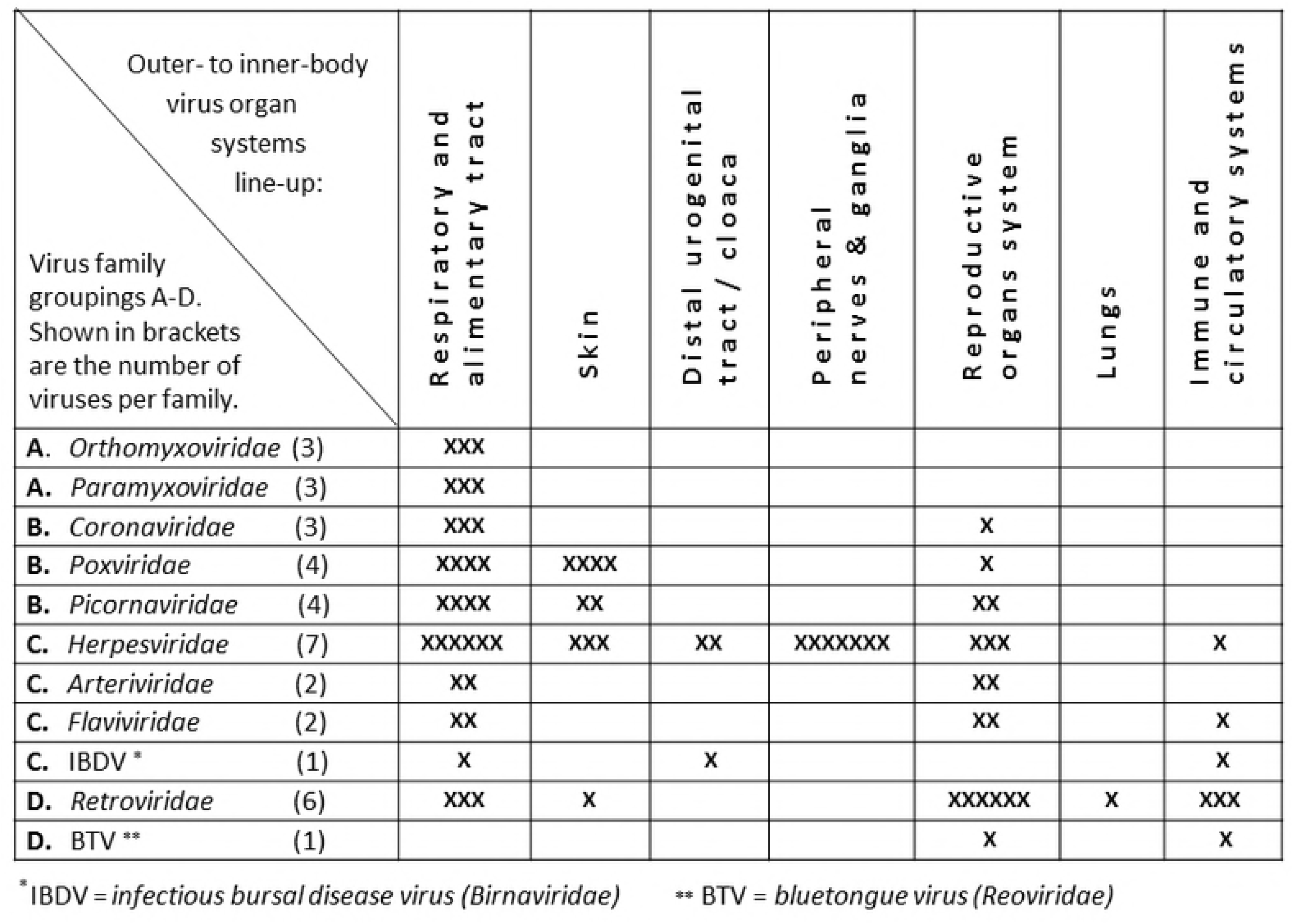
A virtual, outer- to inner-body line-up of organ systems. The organ system tropisms of the viruses belonging to each family were collectively fitted to and with the naked eye aligned with the Fig 1 line-up of virus family groupings and families.

### Virus infiltration ranking

#### Criteria for the infiltration ranking of the 36 viruses

Next, guided by the above findings and by the literature data on the transmission ecology of each virus, see S1 Fig, the 36 viruses were ranked in an outer- to inner-body fashion, strictly on ecological grounds. As shown in Fig 3, starting point in the infiltration ranking was the above established, virtual line-up of organ systems. For each organ system or, as appropriate, combination of organ systems, it was examined how the associated infection-shedding-transmission dynamics matched. Considered were the length of the infection and shedding period, virus transmission modes, virus environmental survival, and the infection severity level. The ranking was accordingly refined.

**Fig 3a.**
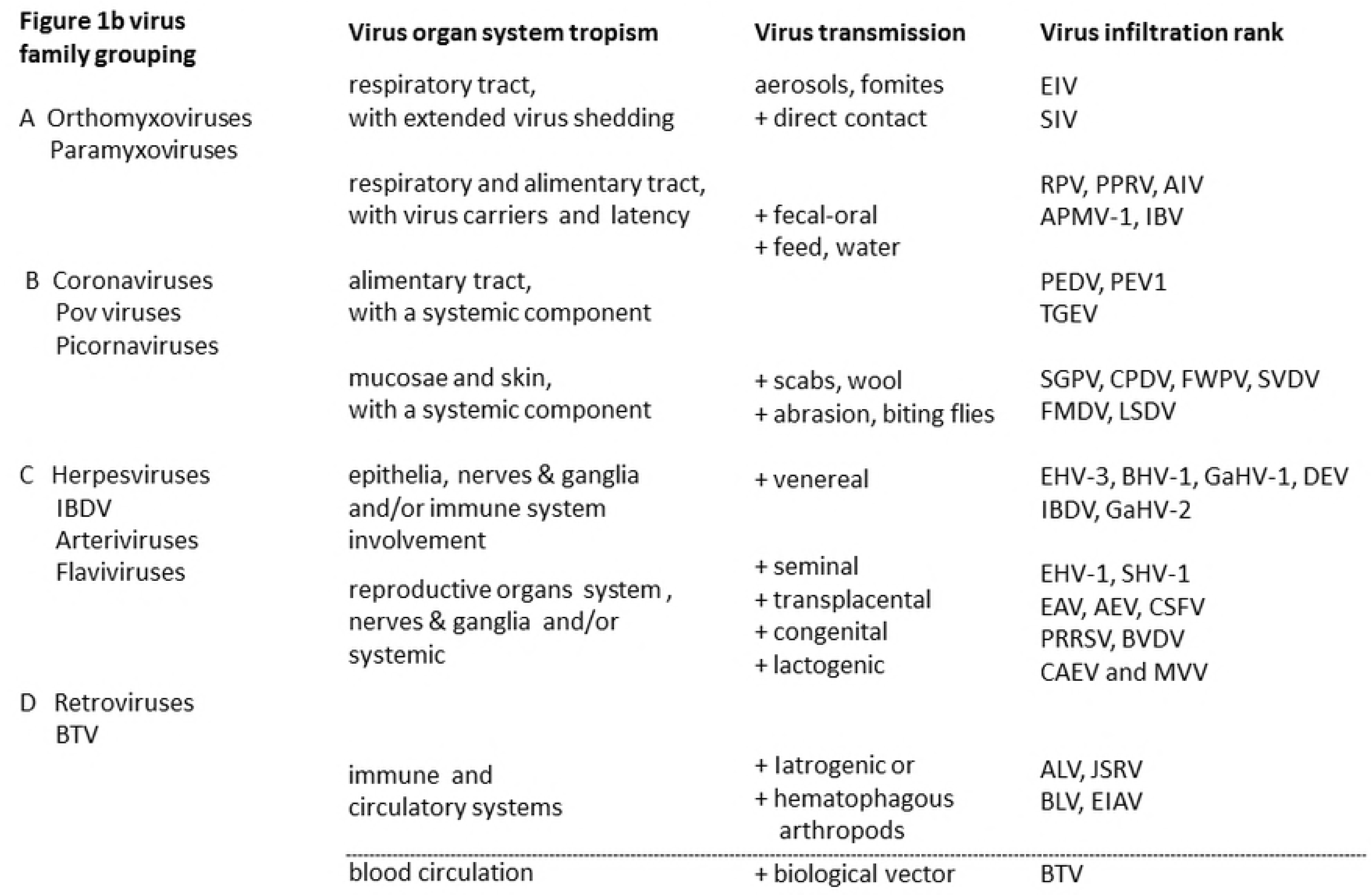
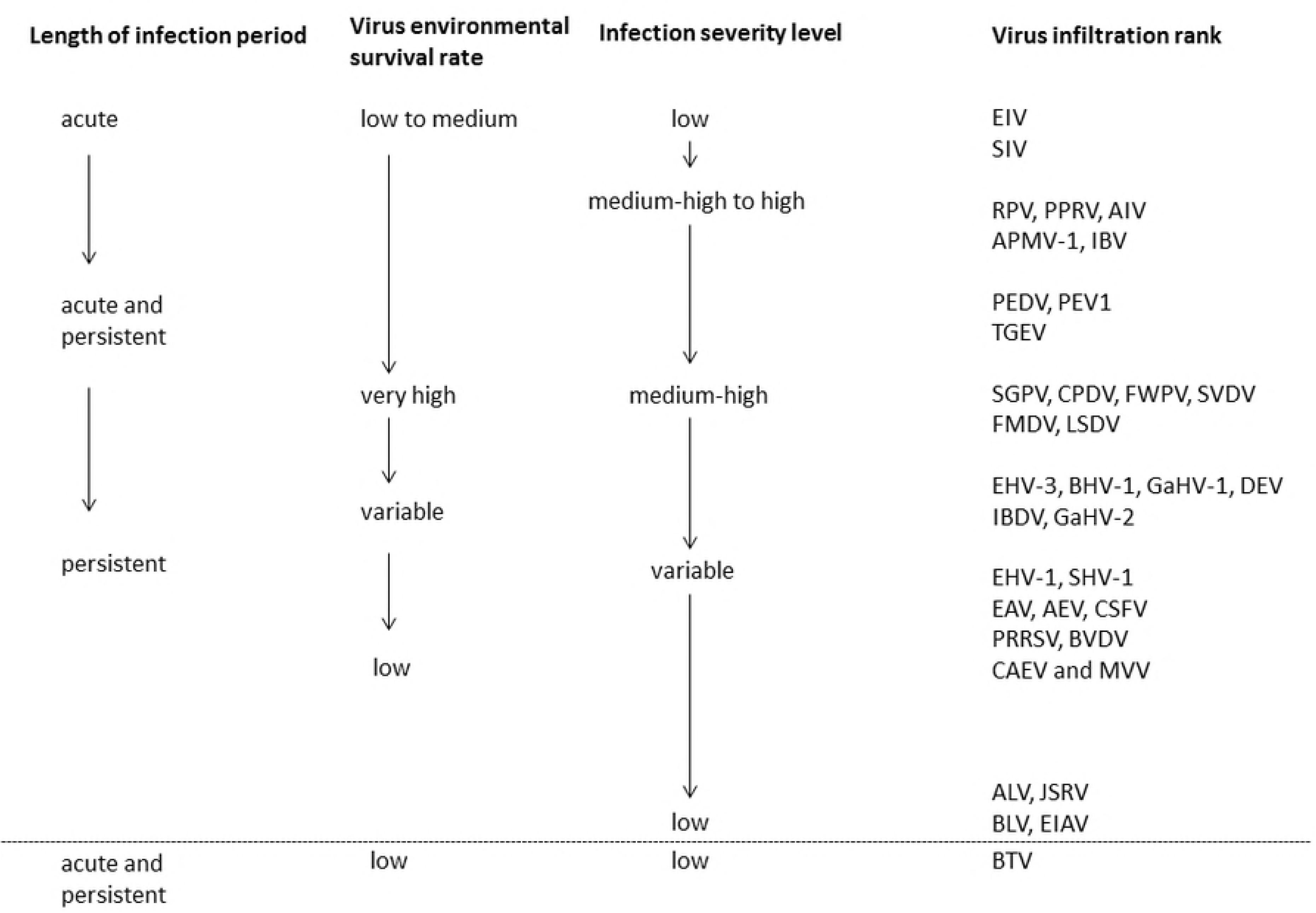
Virus infiltration ranking. The 36 viruses in the study were ranked strictly in ecological terms on the basis of the shifting organ system involvement and associated infection-shedding-transmission dynamics. See Fig 1 for the virus names in full.

#### The ranking of outer-body, epithelial viruses

In brief, from respiratory to alimentary tract to skin, the length of the infection and shedding period and the virus environmental survival rates were found to increase. The viruses colonizing both respiratory and alimentary tract were found to present a distinct, intermediary category. The infection severity level of the latter viruses was found to be relatively high. In general, epithelial viruses shedding over extended periods of time, including after clinical recovery, were considered relatively deep-rooted. Some of the more persistent epithelial viruses featured also non-epithelial transmission modes. Among the skin viruses in the study, the more infiltrative viruses literally invaded all layers of the skin, causing slowly healing lesions. From outside-to-inside, the transmission mode of the pox viruses changed, respectively, from being based on direct plus indirect contact, to direct contact based, involving skin abrasion, or biting flies. With it, the virus environmental robustness tended to decrease.

#### The ranking of inner-body viruses

Infiltration of the inner-body environment was associated with a progressive increase in the share of persistent infections, a decrease in virus environmental robustness, and, very gradually, a reduction in virus pathogenicity. Virus infiltration of epithelia plus peripheral nerves and ganglia, seen for the herpesviruses, typically translated in a persistent or recurrent epithelial infection, not unlike the effects brought away by the immuno-suppressive epithelial viruses in the study. Next, infiltration of the reproductive organs system was found coupled to virus establishment in nerves and ganglia or integral to a systemic infection. The infiltration translated in haphazard virus shedding in semen, abortion during the various stages of pregnancy or, also, orderly virus establishment, as evident from continuous seminal virus shedding, late term abortion, congenital transmission, the combination of congenital and lactogenic transmission, or just lactogenic virus transmission. Next, virus infiltration of the immune and circulatory systems frequently associated with neoplastic disease. For example, infection by the jaagsiekte sheep retrovirus (JSRV) was found to result, albeit in a minority of infections, in the pathognomonic pulmonary adenocarcinoma, supporting virus transmission via virus release in respiratory secretions. Other virus transmission modes comprise iatrogenic transmission and virus transmission involving hematophagous arthropods. The trait profile of the arbovirus in the study, the bluetongue virus (BTV) overlapped with that of the latter category.

#### The virus ranking by organ system

See bottom of the Results section.

#### The interplay of host ecology, virus-host interactions, and virus genome

Finally, as shown in Fig 4, the ranking was disentangled to contrast poultry plus pig and ruminant plus equine viruses, as well as to separately consider the RNA, DNA and retroviruses in the study, ranked in an outer- to inner-body fashion, based on the virus family group line-up shown in Fig 2. Not shown in Fig 2 is the foot and mouth disease virus, a multiple host virus. Spearman correlation revealed the matches among the various virus traits, as pertaining to the two virus host-ecologies, four infection-shedding-transmission related variables, and the three virus genomes.

**Fig 4.**
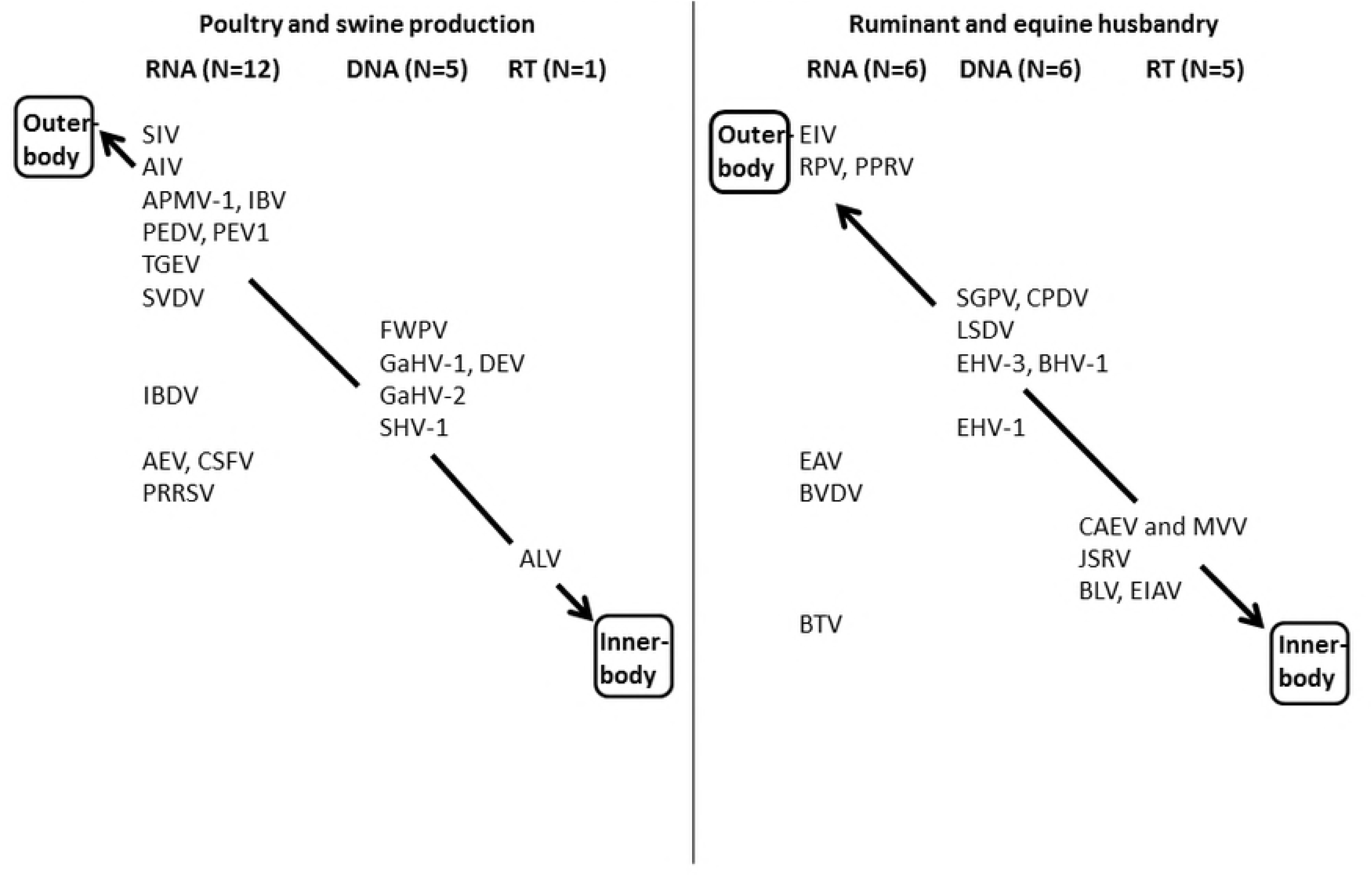
Virus infiltration ranking for two contrasting host ecologies as well as for the RNA, DNA and retroviruses in the study. See text for the matches among virus host-ecology, infection-shedding-transmission variables, and virus genome. See Fig 1 for the virus names in full.

The ruminant and equine viruses turned out to be environmentally less tolerant, with R = - 0.52 and P < 0.005, and causing less severe infections, with R = - 0.48 and P < 0.005. The extent of virus host-body infiltration was found to increase from RNA to DNA to retrovirus, with R = 0.64 and P = 2E-05, for the match with the one-to-four virus infiltration score, and R = 0.50 and P = > 0.005, for the match with the initial, one-to-three score. Also the length of the infection period increased from RNA to DNA to retrovirus, with R = 0.66 and P = 1E-05, while the infection severity level decreased, with R = - 0.39 and P < 0.05. Finally, also the virus genome and the virus host ecology were found correlated. From RNA to DNA to retrovirus, ruminant plus equine viruses increase in prominence, with R = 0.36 and P < 0.05.

### The virus ranking by organ system

#### Respiratory viruses

The equine influenza virus (EIV) was ranked as least infiltrative. In horses, the virus causes a transient respiratory infection. The most important clinical sign is a dry cough usually abating in a few days. Virus transmission is chiefly via aerosols. Fomites may play a role.

Next, in pigs, the swine influenza virus (SIV) causes a respiratory infection involving mucus production and resulting in coughing plus sneezing, aiding virus transmission also on the basis of direct contact. The infection lasts about six days. Virus shedding may continue for up to a month. The virus resists low temperatures; epizootics mainly occur during the winter.

#### Respiratory and alimentary tract viruses

Among the viruses infecting both the respiratory and alimentary tract the rinderpest virus (RPV) was ranked first. Infection in cattle and buffaloes translates in oculonasal discharge, excessive salivation and diarrhea. The virus is typically transmitted during close, muzzle-to-muzzle contact. The virus is mainly excreted during the first week. Fomites are not a viable means of transmission. There are no carriers.

Next, the infection caused by the closely related peste des petits ruminants virus (PPRV) in sheep and goats associates with significant respiratory discharge and diarrhea. The infection takes up to 14 days and there are no carriers. Virus transmission is on the basis of direct contact or via aerosols.

Next, in poultry the avian influenza virus (AIV) causes respiratory and alimentary tract infections, variable in severity. Highly pathogenic avian influenza in chicken usually translates in per-acute death. Mild chicken viruses are also common. Virus transmission is on the basis of aerosols, through direct contact or involving fomites, contaminated feed stuffs or water. The virus is usually excreted for about a week. The virus may survive up to 32 days in the environment.

Next, the avian paramyxovirus type 1 (APMV-1) in poultry is responsible for, respectively, respiratory, gastrointestinal, and nervous signs. In addition, clinically normal birds may carry the virus. Transmission is airborne and fecal-oral. The virus survives well at ambient temperatures in feces.

Next, the infectious bronchitis virus (IBV) in poultry is shed in tracheal mucus and feces. The infection is mostly resolved within 14 days. Also latent infection and erratic virus shedding may occur.

#### Alimentary tract viruses

Among the viruses colonizing the alimentary tract, the porcine epidemic diarrhea virus (PEDV) in pigs typically causes diarrhea. The infection lasts six to 35 days after the onset of clinical signs. Transmission is fecal-oral, involves fomites, feed, or people. The virus survives for weeks outside the body.

Next, the porcine enterovirus encephalomyelitis virus (PEV1) in pigs is excreted in feces and in oral secretions. Recovered animals may continue to shed virus for up to seven weeks. Virus transmission is fecal-oral. The virus may survive in the environment for three months.

Next, the transmissible gastroenteritis virus (TGEV) in pigs causes vomiting and profuse diarrhea, with a systemic component. Mortality is highest in young piglets not protected by colostral antibodies. In adults, recovery takes five to ten days. Carrier sows may shed virus in feces or milk. Transmission is via aerosols, through direct contact, fomites or vertically via the milk. The virus resists low temperatures; epizootics occur mainly during the winter.

#### Mucosal and skin viruses

Among the viruses colonizing mucosae and skin the sheep and goat pox virus (SGPV) in small ruminants was ranked first. The pox lesions take weeks to heal. The clinical signs comprise oculonasal discharge. Virus transmission is direct or indirect. The virus is extremely robust as it may remain infective in scabs for two months and for six months in the environment.

Next, the contagious pustular dermatitis virus (CPDV) in small ruminants causes lesions on mouth, muzzle, udder, and feet. Most infections heal in three to six weeks. Virus transmission is mainly through direct contact. After recovery, virus on wool and hide remains infective for up to one month.

Next, the fowlpox virus (FWPV) in chickens and turkeys causes dry skin lesions or diphteritic, wet nodules in mouth and trachea, resulting in a protracted course of infection. The virus slowly spreads in poultry flocks. The virus transmits during close contact or through skin abrasions; the virus is abundantly present in the skin lesions. Also mechanical transmission by biting flies is possible.

Next, the swine vesicular disease virus (SVDV) in pigs causes vesicles on the feet, lower limbs, and snout. Virus excretion from nose and mouth usually stops within two weeks. In addition, there is evidence for a systemic infection. Virus shedding in feces may last up to three months. The virus spreads mainly through direct contact. Also swill feeding may start an outbreak. The virus is environmentally resistant.

Next, the foot and mouth disease virus (FMDV) in cloven-hoofed animals may during incubation be shed in milk and semen. Upon onset of clinical signs virus may be present in breath, saliva, feces, and urine. Vesicles are found on the feet, buccal mucosa, and teats. The carrier stage may last up to six months. Transmission is via aerosols, through direct contact, fomites, or seminal. In pigs, swill feeding may play a role. The virus is environmentally resistant.

Ranked next was the lumpy skin disease virus (LSDV) in cattle causes pox lesions on skin and mucosae and is shed in milk and semen. Pregnant cows may abort. Virus shedding maybe prolonged as acute infection translates in a carrier stage. Virus transmission is mainly by biting flies, mechanically.

#### Epithelial viruses also latently present in nerves & ganglia and/or immune system

Among the viruses colonizing the epithelia and infiltrating peripheral nerves plus ganglia and/or the immune system, the equine herpesvirus-3 (EHV-3) was ranked first for causing non-invasive lesions on the external genitalia which tend to heal in ten to fourteen days. Transmission is on the basis of sexual or genitonasal contact. The virus is environmentally labile.

Next, the bovine herpesvirus-1 (BHV-1), infectious bovine rhinotracheitis or infectious pustular vulvovaginitis/infectious pustular balanoposthitis virus in cattle and buffalo typically infects the respiratory tract or the external genitalia. In the latter scenario, the virus may be secreted from vagina or prepuce. Virus may also be present in semen. In addition, venereal infection may trigger abortion or a fatal systemic infection in new-borne calves. The venereal infection lasts five to ten days while virus shedding may continue two to sixteen days after infection.

Next, the gallid herpesvirus-1 (GaHV-1) or avian laryngotracheitis virus infection results in the production of bloodstained mucus along the length of the trachea. Virus is latently present in the trigeminal ganglion from where it may reactivate and trigger a recurrent respiratory infection. Virus transmission is via aerosol, through direct contact and fomites. The virus may persist for months in tracheal mucus or carcasses.

Next, the duck enteritis virus (DEV) in ducks, geese and swans is responsible for acute diarrhea. Clinical signs comprise oculonasal discharge. Virus transmission is through direct contact and fomites. Fecal contamination of eggshells may transmit the virus to hatching chicks. Latently infected birds may continue to shed the virus in feces.

Next, the infectious bursal disease virus (IBDV) is an immune-suppressive enteric birnavirus of domestic fowl. The disease is most common in three to six weeks old birds. The virulence level varies. The cloacal bursa is enlarged. The virus is shed in feces. Recovered birds may carry the virus for long periods of time. The virus is environmentally resistant and readily transmitted mechanically among farms by people, equipment and vehicles.

Next, the gallid herpesvirus-2 (GaHV-2) or Marek disease virus in poultry is responsible for visceral lymphomas and neuropathological disorder. Proliferation of lymphoid cells is typically followed by a latency phase, immunosuppression, and formation of neoplasms. Virus is shed from feather follicles for life. Virus transmission results from inhalation of infected dust. The virus remains infective for months to years.

#### Viruses of reproductive organs, also latent in nerves & ganglia and/or systemic

Among the viruses infiltrating the reproductive organs system, peripheral nerves and ganglia, and/or causing a systemic infection, the equine herpesvirus-1 (EHV-1) or equine rhino-pneumonitis virus causes respiratory, neurological and reproductive disorders. The respiratory infection is usually mild and subsides within two weeks. Abortion tends to occur during the third semester. Abortion storms may result from subsequent lateral, airborne virus transmission. Infection around the time of parturation may translate in the birth of a weak, leukopenic foal which dies after a few days.

Next, the swine herpesvirus-1 (SHV-1) or Aujeszky disease virus in pigs may also cause neurological, reproductive, and respiratory disorders. Haphazard abortion, stillbirth and birth of infected piglets occur. Virus reactivation from nerves and ganglia may translate in virus shedding in vaginal secretions, semen and colostrum. Hence, virus transmission is via aerosols, venereal, congenital, and lactogenic. Also swill feeding may kick-start an outbreak; the virus survives up to five weeks in pig meat.

Next, orderly virus establishment in the reproductive organs system as part of a systemic infection is observed for the equine arteritis virus (EAV). The virus usually causes a mild infection. Signs in horses may include nasal discharge, rhinitis, conjunctivitis, edema, dyspnea, diarrhea, and abortion. Stallions tend to become long term carriers and continue to shed virus in semen. Transmission is by aerosols, seminal and, incidentally, congenital.

Next, orderly congenital transmission is observed for the avian encephalitis virus (AEV) in poultry. The virus is neurotropic, enterotropic, and establishes in the proximal genetic tract. Congenitally infected chicks may shed virus in feces and so transmit virus to susceptible chicks. Virus may be shed in feces for several weeks. In adults, carriers may result.

Next, the classical swine fever virus (CSFV) infection in carrier sows may typically result in the birth to weak or also sub-clinically infected piglets. Clinical infections vary from severe per-acute to mild and chronic forms. Virus may be present in oculonasal discharge, blood, feces and semen. Hence, transmission is fecal-oral, through direct contact, fomites, venereal or congenital. Also swill feeding may trigger an outbreak; the virus is moderately fragile.

Next, the porcine reproductive and respiratory syndrome virus (PRRSV) causes respiratory disorders in young piglets and reproductive failure in sows. Virus transmission may be both congenital and lactogenic. In addition, the virus may spread via the respiratory route, through direct contact, swill feeding or venereally. Hence, virus may be present in saliva, feces, urine, semen, and colostrum. Persistently infected sows may abort or, also, give birth to weak or apparently healthy infected piglets.

Next, for the bovine viral diarrhea virus (BVDV) congenital transmission plays a central role in the virus transmission cycle. Virus may also be present in saliva, feces, blood, urine, semen, colostrum, and aborted materials. The infections in cattle vary from subclinical to severe to fatal. Signs may include ulceration of the alimentary mucosa, salivation, respiratory signs, leucopenia, abortion, and congenital anomalies. Acutely infected cattle shed the virus for a short period. Intrauterine infection may result in a persistently infected calf. The virus is environmentally sensitive.

Next, the Maedi-Visna virus (MVV) and the caprine arthritis-encephalitis virus (CAEV) form closely related lentiviruses of sheep and goats and rely on lactogenic virus transmission. Most animal become infected early in life, from drinking infected colostrum or milk. Most infections are subclinical and for life. In sheep, progressive dyspnea or neurological signs may be observed. Encephalomyelitis and poly-arthritis tend to develop in goats. Once clinical signs appear the disease is progressive and usually fatal. Virus may be present in respiratory secretions, feces, semen, colostrum and milk. Virus transmission resulting from direct contact is uncommon; the virus is labile.

#### Viruses of immune and circulatory systems

Among the viruses infiltrating the immune and circulatory systems and epithelia, the avian leukosis virus (ALV) in domestic fowl causes subclinical infection, leukaemia-like, proliferative disease of the hemopoietic system, clonal malignancies of the bursal-dependent lymphoid system, or myeloid leukosis. Clinical signs include weakness, diarrhea, dehydration and emaciation. Chicken with subclinical disease usually shed virus into the albumen or yolk of eggs. Congenitally infected chicks remain viremic for life. Virus may be present feces, saliva, and desquamated skin. Virus survival outside the host body is restricted to a few hours.

Next, the jaagsiekte sheep retrovirus (JSRV) causes clinical signs only in the minority of sheep that develop pulmonary adenomatosis. The virus is more readily detectable in peripheral blood leucocytes and lymphoid organs than in the lungs. Incubation varies from six months to three years. Once a lung tumor develops, the signs are progressive and comprise emaciation, weight loss and respiratory compromise, ending in severe dyspnea and death. Transmission is via milk and colostrum or by the respiratory route, probably via aerosols or droplets. The virus is fragile.

Next, among the fully internalized viruses the bovine leukemia virus (BLV) causes lymphomas in two to five percent of infected animals, predominantly adult cattle older than three to five years. The main transmission is perinatal from cow to calf, via colostrum and milk. Also important are trans-placental transmission, bloodsucking insects, iatrogenic and seminal transmission. The virus is environmentally labile.

Next, the equine infectious anemia virus (EIAV) in horses and other equids may cause a debilitating disease. Subclinical and mild infections are the most common. Bloodsucking flies, in particular of the Tabanidae, transmit the virus mechanically. Also intrauterine and transmission via colostrum or milk occur. The virus is environmentally labile.

#### Virus circulation in the bloodstream

Finally, partially overlapping with the above category are the viruses circulating in the bloodstream and transmitting on the basis of the involvement also of a biological arthropod vector. The bluetongue virus is an arbovirus of sheep plus other domestic and wild ruminants. The virus completes an infection cycle in midges of the genus Culicoides. The virus replicates for six to eight days in the salivary gland after which the midges remain infective for life. In sheep most infections are transient and in-apparent. Seminal and trans-placental transmission may also occur. The epithelial infection does not contribute to the virus transmission success. The virus is environmentally labile.

## Discussion

### A predictive virus evolution framework

A synthesis of the collective findings generated in this study is presented in Fig 5. The host ecology frames the virus transmission modes and, with it, explains the organ system involvement, in turn mirroring the host-cell infection cycle and, with it, the virus genome type. For example, crowding conditions, frequently observed in poultry and pig production, typically attract and select for horizontally transmitting, epithelial viruses. Respiratory and enteric viruses flexibly adjust virulence levels to the dynamic host availability. At the molecular level, these RNA viruses are released from the apical surface of epithelial cells, directly into the outer-body environment. A diametrically opposite scenario is given by the relatively stable host population ecologies observed in ruminant and equine husbandry, with parent stock and their young grazing together in the open, not unlike the herbivores in natural ecologies. The associated viruses count vertically transmitted viruses and also arboviruses. The deep-rootedness of the infection in vital organs systems is at the molecular level matched by virus replication also in non-epithelial cells. Virus latency and low viral replication rates associate with minimal host damage. Retroviruses and other RNA viruses dominate among these ingrained viruses. The DNA viruses in the study, and also the remaining RNA viruses, appear less infiltrative and feature intermediary infection-shedding-transmission and host-cell infection dynamics.

**Fig 5.**
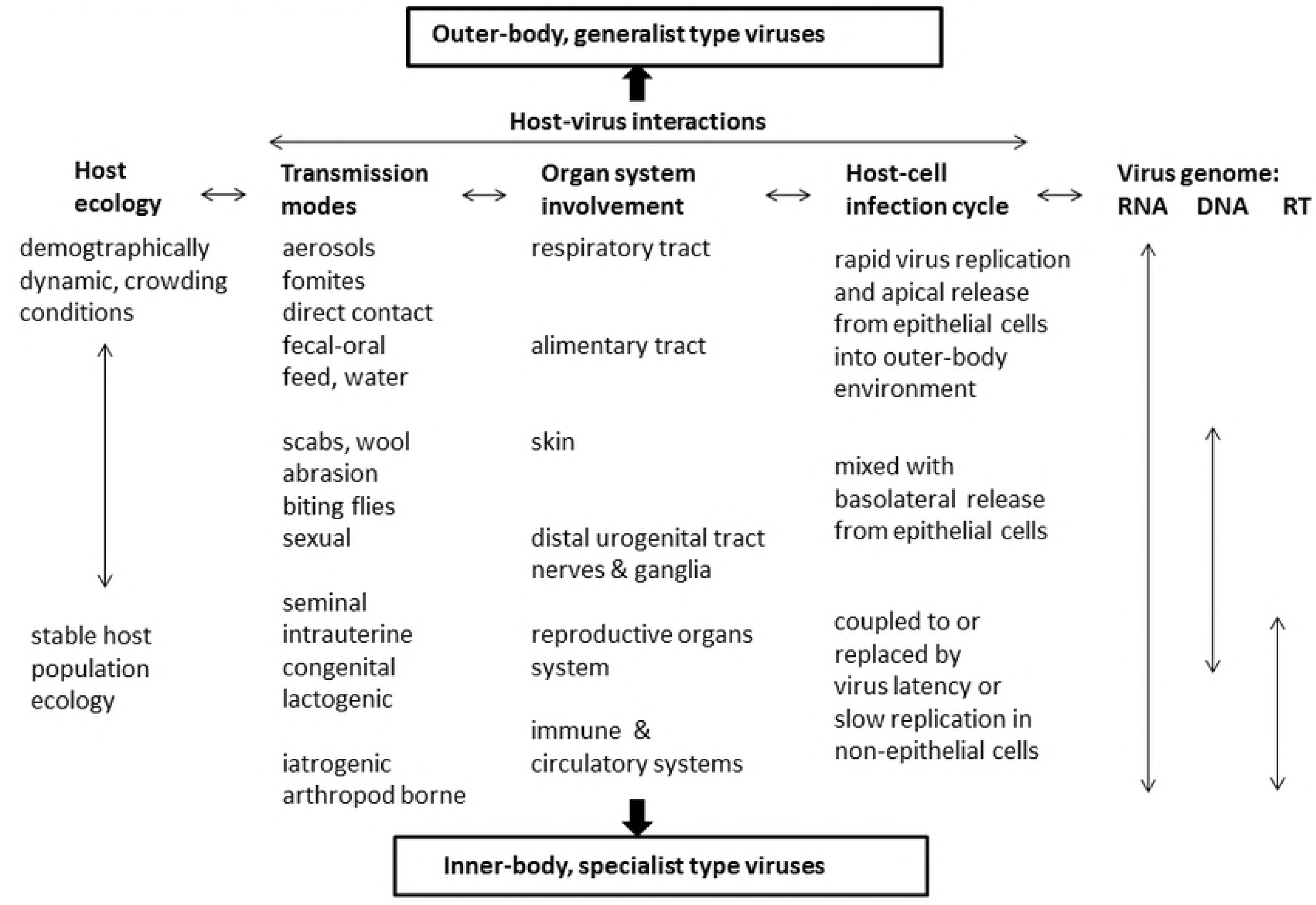
Framework for animal virus ecology and evolution. The framework is based on the outer- to inner-body shifts in the interplay of host ecology, virus-host interactions, and nature of the virus. Two opposing virus evolution pathways emerge.

### Opportunistic viruses and outer-body habitats

Epithelial viruses typically respond to the variation in the environment external to the host-body. Epithelial viruses are highly evolvable [12]. For example, the influenza virus (EIV) in horses generates a transient, dry cough supporting swift virus transmission via aerosols. In pigs, the swine influenza virus (SIV) appears more infiltrative, causing both coughing and sneezing. The latter requires significant mucus production. The infection may last for six days and virus shedding may continue for up to a month. Virus transmission is through direct contact, matching the social behavior of pigs. Likewise, rinderpest virus (RPV) transmits among cattle and buffaloes mainly on the basis of close direct, muzzle-to-muzzle contact. The virus is present in oculonasal discharges and saliva, in addition to being shed in feces. An infection caused by the genetically closely related peste des petits ruminants virus in sheep and goats results in significant respiratory discharge, supporting virus transmission on the basis of direct contact and aerosols. Also the lumpy skin disease virus (LSDV) in cattle and the sheep and goat pox virus (SGPV) circulating in small ruminants are genetically identical. The lumpy skin disease virus causes persistent, deep, necrotic skin plugs and transmits via biting insects, mechanically. The sheep and goat pox virus remains infective in scabs for two months and for six months in the environment. Hence, for epithelial viruses specialization to outer-body virus habitats is integral to virus fitness.

### Specialist viruses and inner-body habitats

Projected on a long evolutionary timescale, the more infiltrative, inner-body viruses tend to become locked in, a consequence of the substitution of the epithelial virus transmission modes by internal organ systems based modes. Virus internalization translates in ever more virus-host intimacy. Least infiltrative among the inner-body viruses are the epithelial herpesviruses latently present also in peripheral nerves and ganglia. Next, virus infiltration of the reproductive organs system may yield vertical transmission modes and so enhance virus-host co-evolution. Next, virus infiltration of the immune and circulatory systems results in more virus-host intimacy still. Eventually, the virus becomes tolerated by the host defense system. Otherwise stated, the virus turns from a parasite into a commensal. Given enough time, the divide between virus and host may become blurred. Also, virus and host may become one [13].

### Health protection implications

Animal virus evolution may be predicted from the position and movement of the virus in the outer- to inner-body gradient. This may be further illustrated with examples of disease emergence from across the human-livestock-wildlife continuum. An opportunistic, epithelial virus of wildlife origin may be found circulating in livestock prior to becoming first detected in humans. This phenomenon is typical for Influenza A viruses [14]. Other examples comprise Henipah viruses [15] and MERS corona virus [16]. SARS corona virus infected civet cats raised as food animals prior to emerging in humans as host [17]. In contrast, inner-body viruses, specialists of vital organ systems, such as the viruses circulating in the bloodstream of non-human primates, may directly jump to humans as host. The driving forces comprise a complex of ecological, socio-economic, demographic and other factors. Examples comprise HIV-aids [18], Chikungunya [19], and Zika viruses [20]. REF Hence, knowing which animal viruses to look for, in farming landscapes and natural ecologies, may assist early warning, early detection of novel viruses and perhaps aid the prevention of novel disease emergence.

## Methods

### Selection of the world main livestock viruses

A subtotal of 23 livestock viruses of major animal health significance was extracted from the OIE-Listed diseases, infections and infestations in force in 2018 [21]. Livestock diseases resulting from virus spill-over from wildlife were excluded from the analysis. The livestock hosts, described in the colloquial OIE terminology, comprise horses, donkeys, cattle, buffaloes, sheep, goats, swine, chicken, turkeys, ducks, and geese. The virus infections and diseases, again in OIE terminology, and in alphabetical order, comprise: Aujeszky disease, avian infectious bronchitis, avian infectious laryngotracheitis, avian influenza, bovine viral diarrhea, bluetongue, caprine arthritis/encephalitis, classical swine fever, equine arteritis, equid herpesvirus-1, equine infectious anemia, equine influenza, enzootic bovine leukemia, foot and mouth disease, infectious bovine rhinotracheitis/pustular vulvovaginitis, infectious bursal disease or Gumboro disease, lumpy skin disease, Newcastle disease, peste des petits ruminants, porcine reproductive and respiratory syndrome, rinderpest, sheep pox and goat pox, and transmissible gastroenteritis. Added to the above were ten globally important livestock infections and diseases drawn from the last, 1995, edition of the Animal Health Yearbook FAO-OIE-WHO [22]: avian encephalitis, avian leukosis, contagious pustular dermatitis, duck virus enteritis or duck plague, enterovirus encephalomyelitis or Teschen disease, fowlpox, Maedi-Visna, Marek’s disease, pulmonary adenomatosis or jaagsiekte, and swine vesicular disease. Furthermore added were three ubiquitous livestock diseases of international animal health significance: equine herpesvirus-3, porcine epidemic diarrhea, and swine influenza, bringing the total to 36 livestock viral infections and diseases. The respective viruses belong to eleven different virus families, and form a mix of RNA, DNA, and retroviruses.

### Literature data on the transmission ecology of the 36 viruses

Shown in S1 Fig are the virus family, the virus genomic architecture, name of the virus, virus abbreviation, and the name given to the infection or disease. In addition, a brief summary is given of the virus transmission ecology, with references to the main livestock host, the virus organ system tropism, the length of the infection and shedding period, the infection severity level, transmission modes, and the virus environmental survival rate.

### Four virus ecological variables

Shown in S2 Fig are one-to-three scores allocated to the 36 viruses for four ecological variables comprising, respectively, the extent of virus host-body infiltration, the length of the infection period, the infection severity level, and the virus environmental survival rate. The one-to-three score for the extent of virus host-body infiltration reflects, respectively, virus transmission resulting from an infection of the epithelia, of epithelia and internal organ systems, or of just internal organ systems. The score for the length of the infection period reflects, respectively, acute, acute and persistent, and persistent infections. Likewise, the score for the infection severity level concerns a case fatality of less than one, one to ten, and above ten percent. The score for the virus environmental survival rate refers to the number of days that the virus remains infective outside the host body, ranging from up to three, three to ten, to over ten days. Also indicated in S2 Fig is the virus host range.

### Literature sources

The literature references, on which the S1 Fig description of the transmission ecology of each virus as well as the virus ecological variable scoring in S2 Fig are based, are detailed in S3 Fig.

### The analysis

The analysis concerned an iterative process. As a first step, the one-to-three scores allocated to the 36 viruses for the four ecological variables were matched on the basis of Spearman correlation.

Next, the eleven virus families in the study were grouped and ranked on the basis of the infiltration scores allocated to the family viruses, creating a novel, one-to-four, family specific virus infiltration score.

Next, the new infiltration score was matched to the scores for the other three variables, for the 36 viruses, and, also, separately for the outer- and the inner-body viruses.

Next, the organ system tropisms of the viruses belonging to each family were collectively fitted to and with naked eye aligned with the above established outer- to inner-body line-up of virus family groups, yielding a virtual outer- to inner-body line-up of organ systems.

As a next step, guided by the above findings and by the literature data on the transmission ecology of each virus, see S1 Fig, the 36 viruses were ranked in an outer- to inner-body fashion, strictly on ecological grounds. Starting point in the ranking was the line-up of organ systems. For each organ system or, as appropriate, combination of organ systems, it was examined how the infection-shedding-transmission dynamics matched. Considered were the length of the infection and shedding periods, virus transmission modes, the infection severity level, and the virus environmental survival rate. The ranking was accordingly refined.

Finally, the above ranking was disentangled to contrast poultry plus pig and ruminant plus equine viruses, as well as to separately consider the RNA, DNA and retroviruses in the study, with the latter ranked in an outer- to inner-body fashion, as indicated by the above established virus family group line-up. Spearman correlation served to reveal the matches among the various virus traits, as pertaining to the two virus host-ecologies, the four infection-shedding-transmission related variables, and the three virus genomes. Viruses circulating in both host species groups were excluded from correlations involving the host ecology.

## Acknowledgment

I am grateful to Epke Le Rütte, Lenny Hogerwerf, Anneke Engering, Marjan Leneman, Jelle Bruinsma, Dorothea van Ooyen and Marleen Slingenbergh for discussions.

## Supporting information

**S1 Figure.**
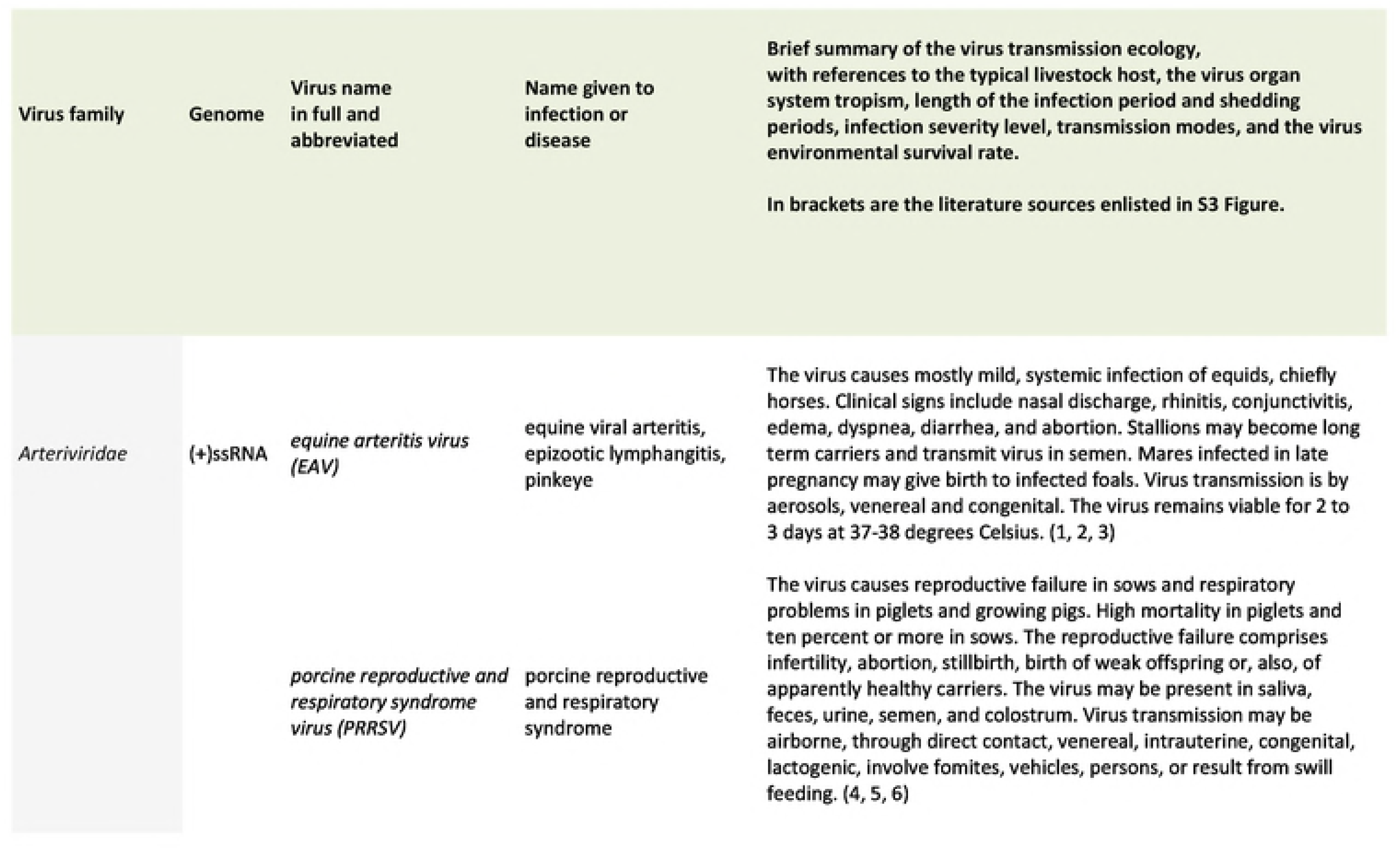
Brief summary of the virus transmission ecology of each of the 36 viruses. References are made to the primary livestock host, the virus organ system tropism, the length of the infection and shedding period, the infection severity level, the transmission modes, and the virus environmental survival rate. Also indicated are references to the literature sources detailed in S3 Fig.

**S2 Figure.**
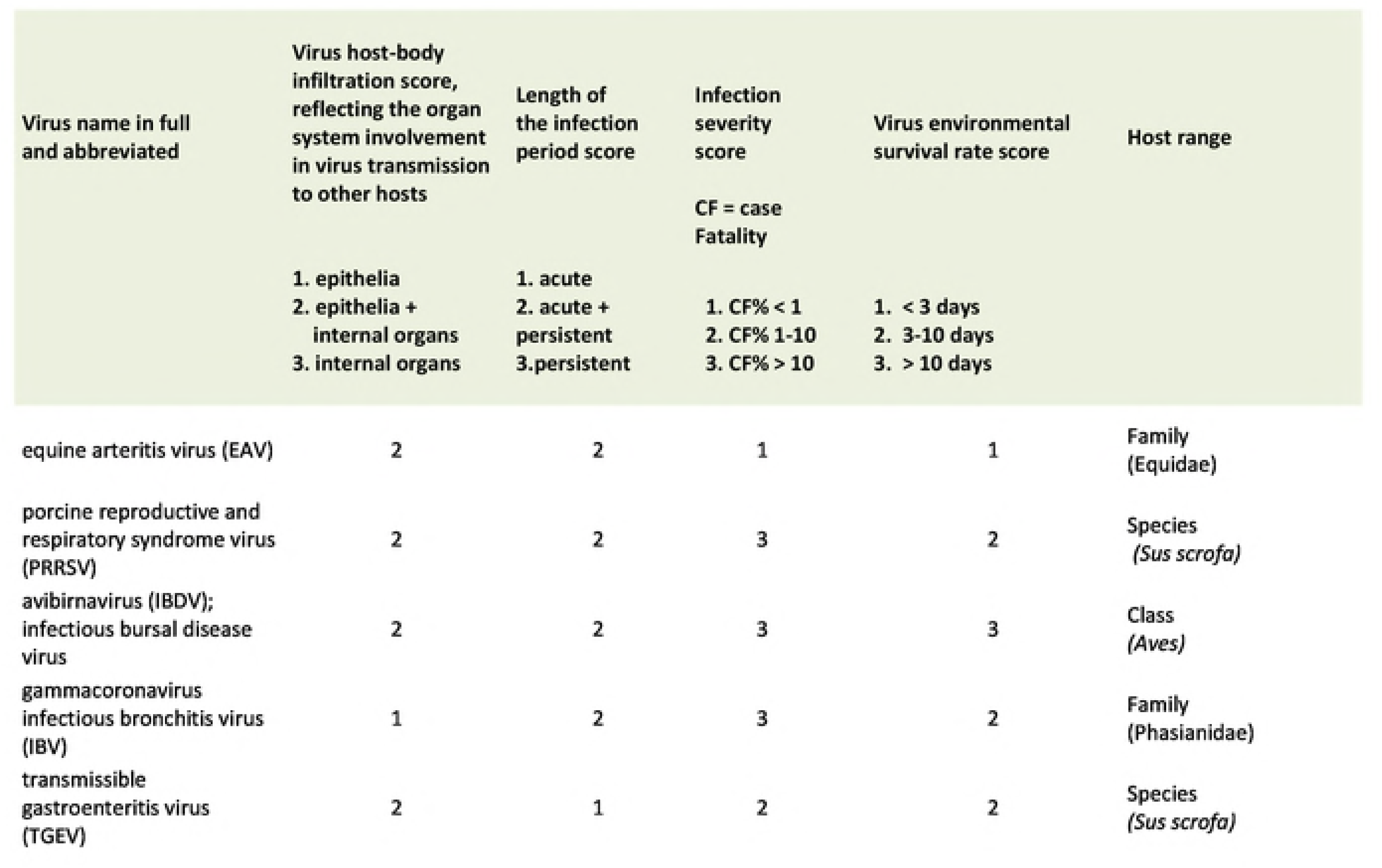
One-to-three scoring of four virus ecological variables. The four virus ecological variables comprise, respectively, the extent of virus host-body infiltration, the length of the infection period, the infection severity level, and the virus environmental survival rate. The one-to-three score for the extent of virus host-body infiltration reflects, respectively, virus transmission resulting from an infection of the epithelia, of epithelia and internal organ systems, or of just internal organ systems. The score for the length of the infection period reflects, respectively, acute, acute and persistent, and persistent infections. Likewise, the score for the infection severity level concerns a case fatality of less than one, one to ten, and above ten percent. The score for the virus environmental survival rate refers to the number of days that the virus remains infective outside the host body, ranging from up to three, three to ten, to over ten days. Also shown are the virus host range and references to the literature sources detailed in S3 Fig.

**S3 Figure.**
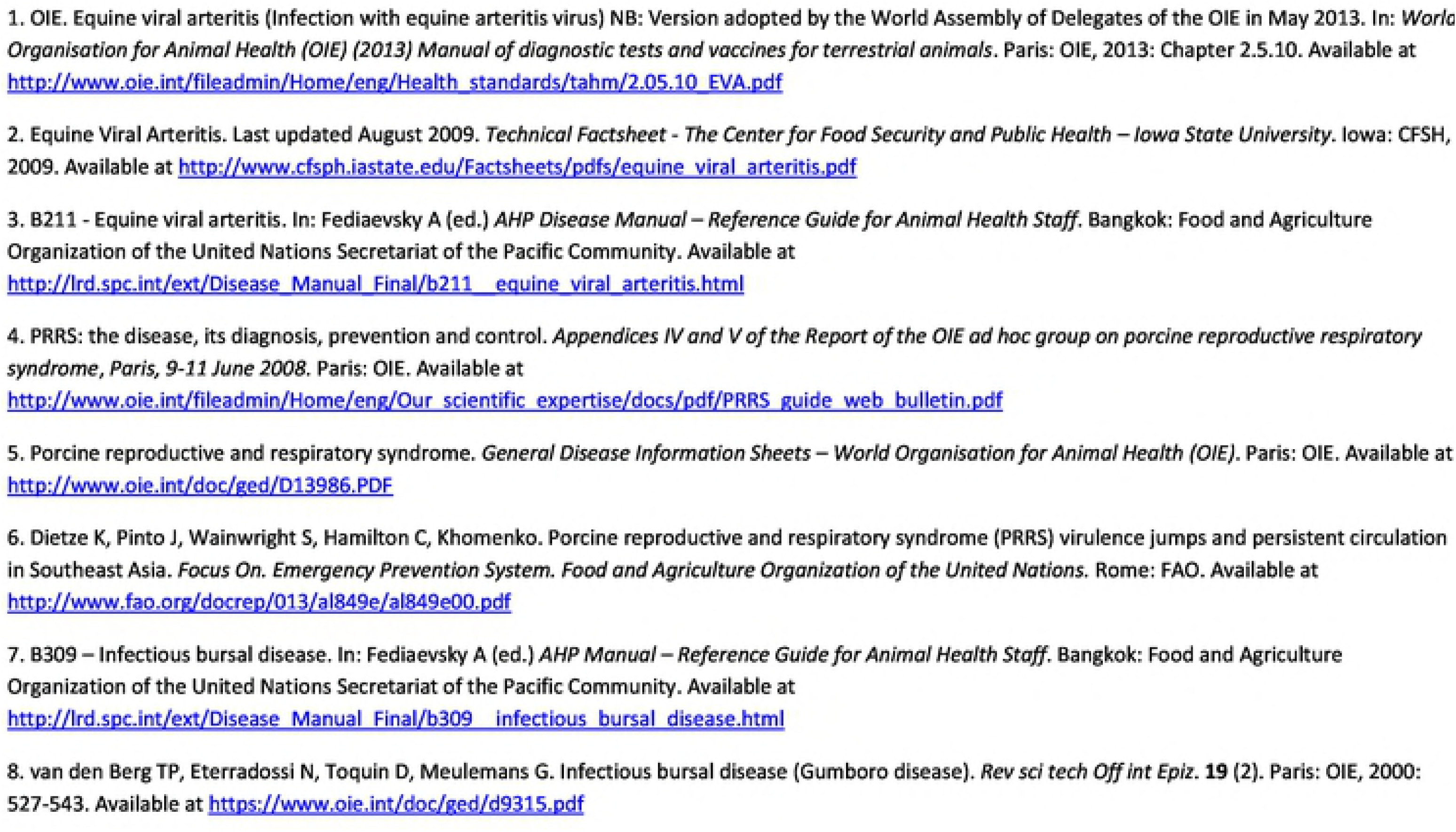
Literature sources for S1 and S2 Figs. The literature references, on which the brief description of the transmission ecology of each of the 36 viruses in S1 Fig as well as the scoring of the four virus ecological variables in S2 Fig are based, are detailed in S3 Fig.

